# Role of spatial patterning of N-protein interactions in SARS-CoV-2 genome packaging

**DOI:** 10.1101/2021.01.06.425605

**Authors:** Ian Seim, Christine A. Roden, Amy S. Gladfelter

## Abstract

Viruses must efficiently and specifically package their genomes while excluding cellular nucleic acids and viral sub-genomic fragments. Some viruses use specific packaging signals, which are conserved sequence/structure motifs present only in the full-length genome. Recent work has shown that viral proteins important for packaging can undergo liquid-liquid phase separation (LLPS), where one or two viral nucleic acid binding proteins condense with the genome. The compositional simplicity of viral components lends itself well to theoretical modeling compared to more complex cellular organelles. Viral LLPS can be limited to one or two viral proteins and a single genome that is enriched in LLPS-promoting features. In our previous study, we observed that LLPS-promoting sequences of SARS-CoV-2 are located at the 5ʹ and 3ʹ ends of the genome, whereas the middle of the genome is predicted to consist mostly of solubilizing elements. Is this arrangement sufficient to drive single genome packaging, genome compaction, and genome cyclization? We addressed these questions using a coarse-grained polymer model, LASSI, to study the LLPS of nucleocapsid protein with RNA sequences that either promote LLPS or solubilization. With respect to genome cyclization, we find the most optimal arrangement restricts LLPS-promoting elements to the 5ʹ and 3ʹ ends of the genome, consistent with the native spatial patterning. Genome compaction is enhanced by clustered LLPS-promoting binding sites, while single genome packaging is most efficient when binding sites are distributed throughout the genome. These results suggest that many and variably positioned LLPS-promoting signals can support packaging in the absence of a singular packaging signal which argues against necessity of such a feature. We hypothesize that this model should be generalizable to multiple viruses as well as cellular organelles like paraspeckles, which enrich specific, long RNA sequences in a defined arrangement.

**Statement of significance:** The COVID-19 pandemic has motivated research of the basic mechanisms of coronavirus replication. A major challenge faced by viruses such as SARS-CoV-2 is the selective packaging of a large genome in a relatively small capsid while excluding host and sub-genomic nucleic acids. Genomic RNA of SARS-CoV-2 can condense with the Nucleocapsid (N-protein), a structural protein component critical for packaging of many viruses. Notably, certain regions of the genomic RNA drive condensation of N-protein while other regions solubilize it. Here, we explore how the spatial patterning of these opposing elements promotes single genome compaction, packaging, and cyclization. This model informs future *in silico* experiments addressing spatial patterning of genomic features that are experimentally intractable because of the length of the genome.

## Introduction

Biomolecular condensation is a simple and versatile way for cells to spatially and temporally control biochemistry. It is now clear that a wide variety of compartments likely form using the process of liquid-liquid phase separation (LLPS) which leads to a condensation of specific components out of bulk cytosol or nucleoplasm (1). The protein components of condensates tend to contain intrinsically disordered or low complexity sequences and RNA-binding domains (2). Many condensates also contain nucleic acids, and indeed RNA can promote phase separation in many instances (3). However, the contributions of specific RNA sequences and structures in condensate assembly, contents, and material properties is poorly understood (3).

Viruses present a powerful system to examine sequence specificity for both proteins and nucleic acids in phase separation because of their highly compact genomes and limited protein coding genes. Indeed, reports have emerged for VSV (4), respiratory syncytial virus (5), rabies (6), measles (7), and HIV (8) components showing the capacity to undergo LLPS.

SARS-CoV-2 is a positive strand RNA virus that has an exceptionally large genome of ∼30kb which is selectively packaged into a relatively small capsid estimated to be ∼100 nm in diameter (9). How the genome is selectively packaged while excluding sub-genomic RNAs generated by the virus and the host transcriptome and sufficiently compressed to fit into a virion is not yet understood. The necessity of a packaging signal for SARS-CoV-1 is still not clear, although one sequence has been found to be sufficient but not necessary to package RNA (10). To our knowledge, no packaging signal has been identified for SARS-CoV-2.

The nucleocapsid protein of SARS-CoV-2 undergoes LLPS (11–15), and our work found that this occurs in an RNA sequence-specific manner with different regions of the genome (11). Remarkably, RNAs of the same length can either promote or limit phase separation depending on the sequences. The sequences with differing behavior also show distinct patterns of binding of N-protein with LLPS-promoting sequences having discrete patterned N-protein interactions, while RNA sequences that limit phase separation are uniformly coated in N-protein. The regions that promote phase separation are in the 5ʹ and 3ʹ ends of the genome, prompting us to speculate that phase separation could be relevant to packaging, as these LLPS-promoting sequences are present specifically on the whole genome and would not be together on subgenomic or host RNAs.

Here, we developed a coarse-grained model to test the hypothesis that phase separation could be a relevant process for selecting and compacting a single genome. Our goal was to examine how the linear location of different RNA sequences in the genome generates spatially segregated and condensed RNA molecules. We first explored fragments of the SARS-CoV-2 genome that have opposing phase behavior when mixed with N-protein, as shown in (11). Specifically, the 5ʹ and 3ʹ ends of the genome promote phase separation, while the frameshifting element (FE) and central regions of the genome solubilize N-protein (**Fig 1A**). We next examined the spatial patterning of these opposing elements within a full genome model and quantified its effects on phase separation, packaging of single genomes, genome compaction, and genome cyclization. We found that in this model, localization of LLPS-promoting features to the 5ʹ and 3ʹ ends of the genome is sufficient to drive LLPS-based single genome packaging and genome compaction, and is necessary for genome cyclization. Addition of clustered LLPS-promoting features throughout the genome further enhanced all of these metrics.

**Figure 1:**
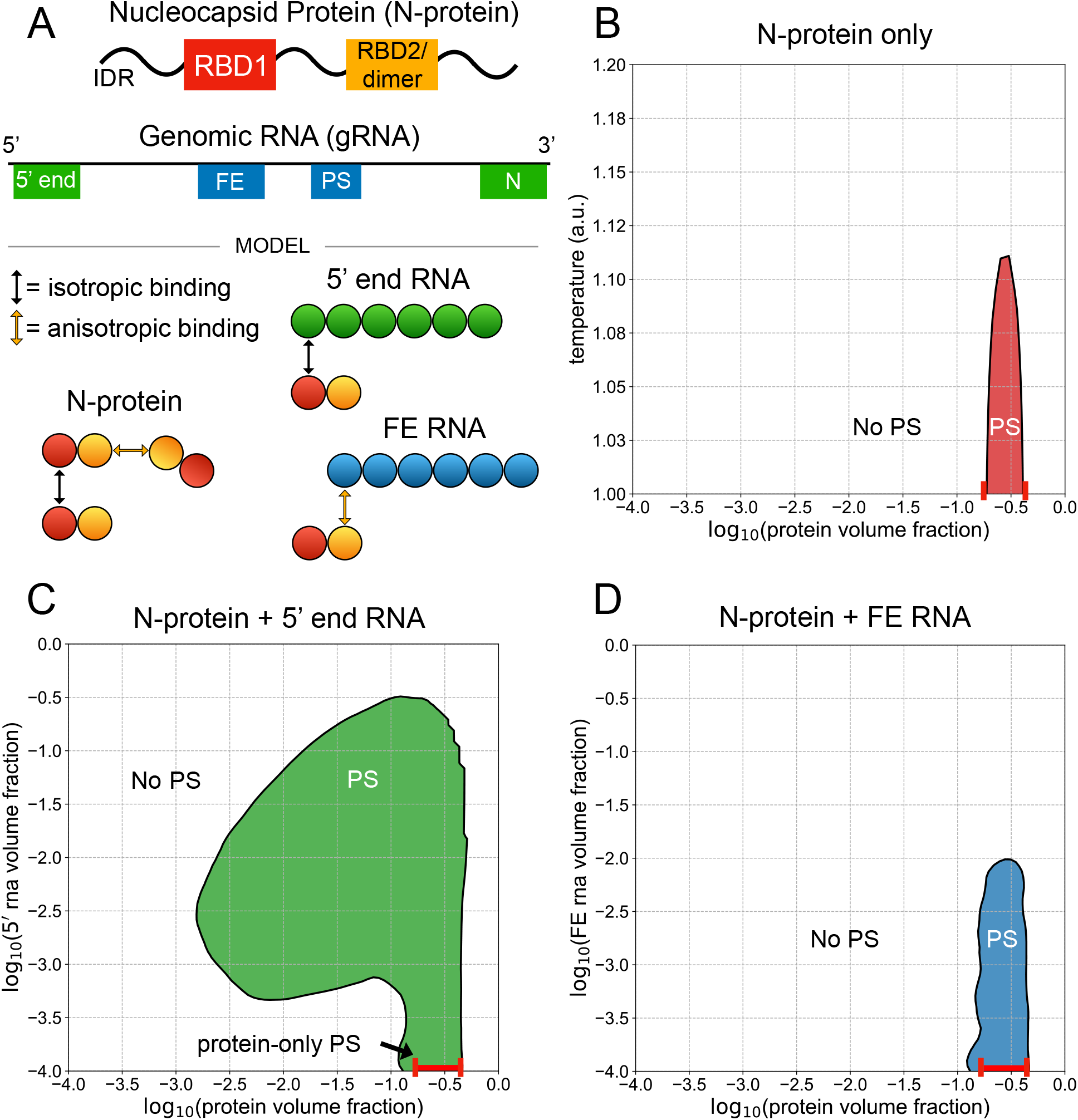
5ʹ end and FE RNA with N-protein have opposing phase behavior. (A) N-protein is represented as a two-bead chain, with the first bead participating in isotropic homotypic interactions, and the second bead participating in anisotropic homotypic interactions. Both 5ʹ end and FE RNA segments are roughly three times larger than N-protein and are represented by six beads each. N-protein interacts with all 5ʹ end RNA beads via isotropic binding with its first bead, and it interacts with all FE RNA beads via anisotropic binding with its second bead. This interaction with FE competes with N-protein dimerization. (B) N-protein phase separates (PS) in a narrow concentration and temperature range on its own. (C) N-protein with 5ʹ end RNA at temperature 1 a.u. phase separates across a wider concentration range than on its own. (D) Nprotein with FE RNA at temperature 1 a.u. is solubilized at sufficiently high FE RNA concentrations.

## Methods

### LASSI model parameterization

Simulations were performed using LASSI (16) and run on the Longleaf computer cluster at UNC-Chapel Hill and on the Comet XSEDE cluster at the San Diego Supercomputer Center (17). Each simulation was run independently on a single compute node with 4GB RAM. The following default parameter sets were used for all simulations (Tables 1, 2):

**Table 1:** Isotropic binding energies.

| **Isotropic Binding Energies** | RBD1 | RBD2/dimer | 5' end RNA | FE RNA | 5' gRNA terminus | 3' gRNA terminus |
| --- | --- | --- | --- | --- | --- | --- |
| RBD1 | -0.5 | 0.0 | -1.2 | 0.0 | 0.0 | 0.0 |
| RBD2/dimer | 0.0 | 0.0 | 0.0 | 0.0 | 0.0 | 0.0 |
| 5' end RNA | -1.2 | 0.0 | 1.0 | 1.0 | 1.0 | 1.0 |
| FE RNA | 0.0 | 0.0 | 1.0 | 1.0 | 1.0 | 1.0 |
| 5' gRNA terminus | 0.0 | 0.0 | 1.0 | 1.0 | 1.0 | 0.0 |
| 3' gRNA terminus | 0.0 | 0.0 | 1.0 | 1.0 | 0.0 | 1.0 |

**Table 2:** Anisotropic binding energies.

| **Anisotropic Binding Energies** | RBD1 | RBD2/dimer | 5' end RNA | FE RNA | 5' gRNA terminus | 3' gRNA terminus |
| --- | --- | --- | --- | --- | --- | --- |
| RBD1 | 0.0 | 0.0 | 0.0 | 0.0 | 0.0 | 0.0 |
| RBD2/dimer | 0.0 | -3.0 | 0.0 | -5.0 | 0.0 | 0.0 |
| 5' end RNA | 0.0 | 0.0 | 0.0 | 0.0 | 0.0 | 0.0 |
| FE RNA | 0.0 | -5.0 | 0.0 | 0.0 | 0.0 | 0.0 |
| 5' gRNA terminus | 0.0 | 0.0 | 0.0 | 0.0 | 0.0 | -3.0 |
| 3' gRNA terminus | 0.0 | 0.0 | 0.0 | 0.0 | -3.0 | 0.0 |

Negative energies indicate attraction and positive energies indicate repulsion. The appropriate subsets of these interactions were used for the simulations involving N-protein alone, N-protein and 5ʹ end RNA, N-protein and FE RNA, and all spatial rearrangements of beads in genomic RNA (gRNA) simulations. We performed parameter sweeps over many of these energies for the Nprotein only system, the 5ʹ end and FE RNA fragments with N-protein systems, and the Nprotein with gRNA systems, explained further in the main text and SI (**Figs S2-S4, S7, S8**). Simulations of N-protein alone were run at 15 temperatures linearly spaced between 1 arbitrary unit (a.u.) and 2 a.u., and 15 concentrations logarithmically spaced between 1e-4 and 1e-0.3, each for 1e9 total time steps with 5e6 timesteps of thermalization. Temperature scales interaction energies, *ε*, as 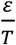. During thermalization, the temperature is raised to 1000 a.u., anisotropic interactions are inactivated, and all chains are pushed to the center of the system, as described in (16). For systems with short chains, like the 6- and 2-bead chains representing 5ʹ end and FE RNA fragments and N-protein, respectively, this step allows for a more rapid convergence to equilibrium if the system will phase separate. For very long chains, this step may kinetically trap the system in a clustered state. For systems with N-protein and 5ʹ end and FE RNA fragments, simulations were run at a single temperature, 1 a.u., for 1e9 timesteps and 5e6 steps of thermalization. For genomic RNA (gRNA) with N-protein systems, simulations were run at a single temperature, 1 a.u., for 10e9 timesteps with no thermalization due to kinetic trapping when thermalization was used. These simulations equilibrated to similar minimum energies with or without thermalization, but had altered packaging properties (**Fig S5, S6**). We concluded that since gRNA chains in thermalized systems gradually disengaged from the initial central cluster over the course of simulations that their equilibrium states were not fully clustered at the center, as observed for simulations without thermalization (**Fig S5**) and that they were instead kinetically trapped. Thus, we did not use thermalization for gRNA systems. For all ternary systems, concentrations of each component were chosen to be logarithmically spaced across the ranges of interest, with hundreds of ([protein], [RNA]) coordinates sampled for each system (**Fig S1**). For N-protein with 5ʹ end and FE RNA, the maximum number of Nprotein and RNA chains were used such that the total beads in the system never exceeded 40000 and the target stoichiometries were satisfied. For simulations with gRNA and N-protein, 10 gRNA molecules were always used, and the number of N-protein chains was altered to match target stoichiometries. At the lowest gRNA and highest N-protein volume fractions, the number of gRNA molecules was gradually scaled down to 1 due to computational limitations on the sizes of the systems. At each coordinate, 2 independent simulations were run with and without interactions, for a total of 4 simulations at each concentration coordinate.

### Simulation analysis

Analysis was performed using scripts within LASSI and custom scripts that relied upon the ovito python module (18). LASSI outputs a global density inhomogeneity value, 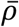, for each simulation, which is used to determine whether phase separation has occurred. 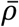 is calculated using the pair distribution function for all beads in the system by default, as described in (16). We used this default calculation for N-protein-only systems and the 5ʹ end and FE RNA fragments with N-protein systems. However, for gRNA systems, we did not observe a dilute phase of gRNA chains and concluded that phase separation of gRNA in this context was not meaningful (more detail in main text). Instead, we calculated a new density inhomogeneity metric, 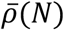, that uses the pair distribution function for only N-protein beads in the system. Using this limited pair distribution function, we calculated 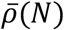) analogously to 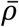 as described in (16). Contours at 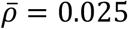 were used to determine phase boundaries for N-protein only and 5ʹ end and FE RNA fragments systems. This value of (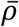was shown in (16) to universally indicate the onset of phase separation. We used 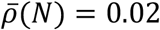 for gRNA systems as this value better aligned with sharp transitions in 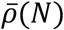 as a function of volume fraction and resulted in smoother phase boundaries. We used the ovito module to calculate clusters using a maximum cutoff of 3^1/2^, the maximum distance between two interacting particles in a cubic lattice. The radius of gyration for each gRNA molecule was calculated according to the equation in the text. Ovito was also used to calculate end-to-end distances of genomic RNAs. We counted genomes as cyclized if their terminal beads were within 3^1/2^ units of each other. For all analysis of packaging metrics in gRNA systems, 100 simulation snapshots were used from the last half of simulations for each of 2 runs and averaged over time and runs. Snapshots were recorded starting at 5e9 steps every 5e7 steps and used for analysis. For 5ʹ end and FE RNA fragments with N-protein systems, 50 simulations snapshots were recorded starting 2e7 steps after thermalization completion and used for analysis. For all systems,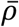 or 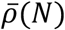 and clustering and packaging metrics were interpolated to a grid linearly spaced between 1e-4 and 1 with a discretization of 80 points along the protein and RNA axes. Interpolation was performed using interpolate.griddata from scipy. Plots were made using matplotlib.

## Results

### Simulations recapitulate known phase behavior and binding patterns of N-protein with FE and 5ʹ end RNA

We first sought to characterize the phase behavior of different regions of the viral genome by focusing on simulations involving the N-protein with either the first 1000 nucleotides of the genome (5ʹ end RNA) or 1000 nucleotides around the frameshifting region located between ORF1A and ORF1B (FE RNA). Simulations were carried out using LASSI, which employs stickersand-spacers representations of polymers and generates full phase diagrams by performing Monte Carlo simulations at many temperatures and volume fractions of components (16). Simulations take place on a cubic lattice, and only a single particle can occupy a given lattice site at one time. In this work, polymer beads are connected by implicit linkers that do not occupy space but rather guarantee that adjacent beads within chains are always in adjacent lattice sites. The first consideration was how to represent the N-protein and RNA in a coarsegrained manner based on existing data. The N-protein has two RNA binding domains (RBDs), a dimerization domain which overlaps with RBD2, and three intrinsically disordered regions (IDRs) (**Fig 1A**). RBD1 is conserved across multiple betacoronavirus genomes and has previously been shown to interact with the conserved sequences and structures in the 5ʹ UTR (19). We previously demonstrated that a single point mutant within RBD1, Y109A, greatly reduces Nprotein phase separation and changes N-protein interactions with 5ʹ end RNA, while only minimally affecting protein binding and phase behavior with FE RNA (11). Since FE RNA primarily solubilized N-protein, and deletion of the RBD2/dimerization domain blocked Nprotein co-phase separation with RNA (12), we postulated that FE may block N-protein phase separation by specifically interacting with RBD2 and preventing N-protein dimerization. Thus, we hypothesize that N-protein RBD1 primarily binds to 5ʹ end RNA, while RBD2 primarily binds to FE RNA.

Based on these data, we represented the N-protein as simply as possible using two spheres. The first sphere participates in weak isotropic interactions with other N-proteins (representing the association of the IDRs) and with the 5ʹ end RNA (representing RBD1). The second sphere participates in strong anisotropic interactions with other N-proteins to capture dimerization and the interaction with the FE RNA via RBD2 (**Fig 1A**). These latter anisotropic interactions operate under the assumption that N-protein dimerization competes with binding to FE RNA, since anisotropic binding in this model is one-to-one. To model charge effects, the RNA molecules experience an isotropic repulsive force amongst themselves, both within chains and among distinct RNA chains. Using these specifications, we sought to qualitatively reproduce the phase behavior among these molecules established experimentally in (11). We found that, relative to N-protein phase separation on its own (**Fig 1B**), addition of 5ʹ end RNA promotes enhanced phase separation across a wide range of protein and RNA concentrations (**Fig 1C**), while addition of FE RNA does not promote phase separation and solubilizes N-protein at sufficiently high concentrations of RNA (**Fig 1D**).

In addition to opposing phase behavior, the binding of N-protein to 5ʹ end and FE RNA as a function of N-protein concentration was shown to be distinctly patterned based on protein crosslinking in (11). While FE RNA is uniformly coated with protein across a wide range of protein concentrations, 5ʹ end RNA has a few discrete binding sites and is only gradually coated more with protein as the protein concentration is increased. The simulations report a similar distinct pattern of protein interactions. For a fixed RNA concentration and at low bulk protein concentrations, 5ʹ end RNA is initially coated with very little protein (**Fig 2A**). As the bulk protein concentration is increased, 5ʹ end RNA sharply transitions to binding large amounts of protein (**Fig 2A**). In contrast, FE RNA binds more protein at low protein concentrations, relative to 5ʹ end RNA, and experiences a more gradual transition to high amounts of bound protein (**Fig 2B**). In all, the distinct protein binding behavior of these two RNA elements is consistent with that found in (11) (**Fig 2C**). Thus, the simulation results are consistent with the experimental system and support that the coarse-grained representations and binding energies are reasonable approximations of the actual system.

**Figure 2:**
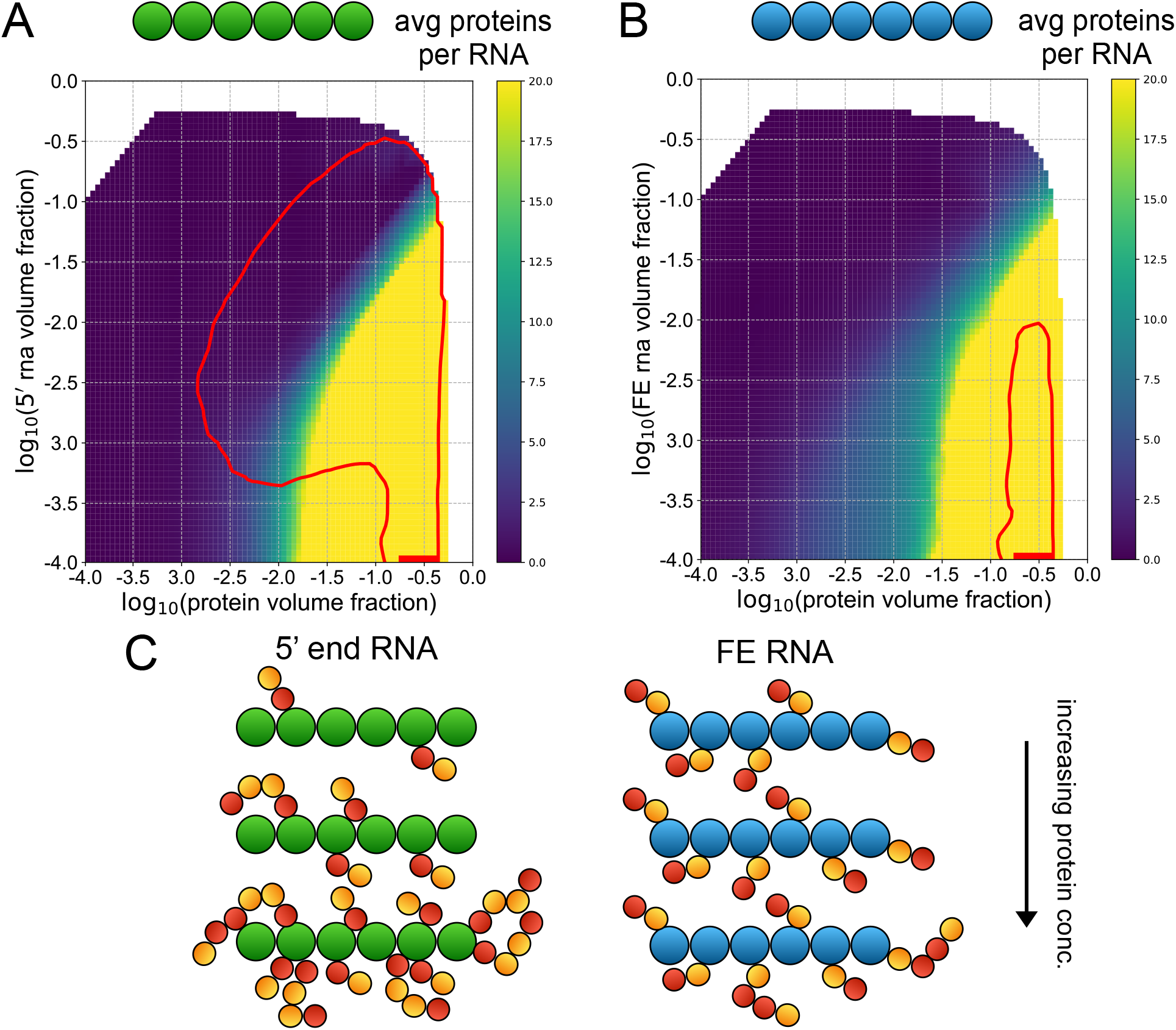
5ʹ end and FE RNA have distinct N-protein binding behavior. (A) and (B) Phase boundaries for 5ʹ end RNA with N-protein (A) and FE RNA (B) with N-protein are shown in red. The heatmaps indicate the average number of protein chains per RNA chain for each cluster identified in each simulation. If more than one RNA chain is in a cluster, the ratio of protein chains to RNA chains within that cluster is reported. (C) For any given fixed RNA concentration, as the protein concentration is increased, 5ʹ end RNA shows a sharp transition to highly bound protein, while FE shows a more gradual and transition, with more proteins bound at low protein concentrations relative to the 5ʹ end system.

We ran extensive parameter sweeps for these systems to determine the contributions of specific interactions. For N-protein alone, the isotropic binding energy greatly influenced phase behavior (**Fig S2**). Dimerization was necessary for phase separation for low isotropic binding energies, including the default energy used for all following simulations (**Fig S2**). Interestingly, the strength of the dimerization interaction was non-monotonically related to phase separation propensity, with very high dimerization energies starting to weaken phase separation (**Fig S2**). We also studied the effects of altering the strength of binding between N-protein and 5ʹ end and FE RNA. The strength of the isotropic binding interaction between N-protein and 5ʹ end RNA strongly influenced co-phase behavior (**Fig S3**). In contrast, the strength of the anisotropic binding between N-protein and FE RNA did not alter phase behavior until very high energies, when a new arm of the phase diagram emerged (**Fig S3**). We did observe differences in the average number of N-proteins bound to FE outside of the phase envelope, with low anisotropic energies leading to less binding (**Fig S3**). We doubled the magnitude of the RNA-RNA repulsive isotropic energy and observed small effects on the phase behavior of N-protein with both 5ʹ end RNA and FE RNA (**Fig S4**). Finally, we increased the isotropic binding energy between Nproteins and observed phase behavior in the context of 5ʹ end and FE RNA. Since N-protein alone phase separates across a wider range of volume fractions under this condition, we see a corresponding widening of the phase diagrams across these concentrations when mixed with both RNA fragments (**Fig S4**). Additionally, more FE RNA is required to solubilize N-protein since the N-proteins are more stably phase separated (**Fig S4**). The most sensitive interactions for these systems appear to be the isotropic binding between N-proteins and the isotropic binding of N-protein with 5ʹ end RNA.

### Whole genome simulations reveal effects of phase separation on single genome packaging, genome compaction, and genome cyclization

The large size (30kb) of the genome makes it challenging to synthesize *in vitro* for experiments. We therefore were eager to use this simulation space to ask questions about how the different RNA-sequence elements will behave when present *in cis* on the same polymer, as they are found in the native virus. To address if specific arrangements of RNA encoded features could be sufficient for packaging a single genome, we utilized the same representations of the 5ʹ end and FE RNA and the N-protein described above and assembled a system that represents the N-protein and the entire viral genome (gRNA) (**Fig 3A**). In addition to the 5ʹ end of the gRNA, the 3ʹ end was also found to promote phase separation with N-protein and shared similar protein binding behavior and RNA sequence features (**Fig 1A**) (11). Central regions of the gRNA that were studied behaved similarly to the FE RNA, and further, the internal portions of the genome were predicted to be more similar to the FE than 5ʹ end RNA (11). Taking into account the relative sizes of nucleotides and amino acids, the gRNA is roughly 90x larger than N-protein, so our representation involves 180-bead chains for each genome. An additional strong, anisotropic interaction between the terminal beads of each gRNA chain is added to represent known nucleotide complementarity between the 5ʹ and 3ʹ ends of the gRNA and propensity to crosslink *in vivo* (20).

**Figure 3:**
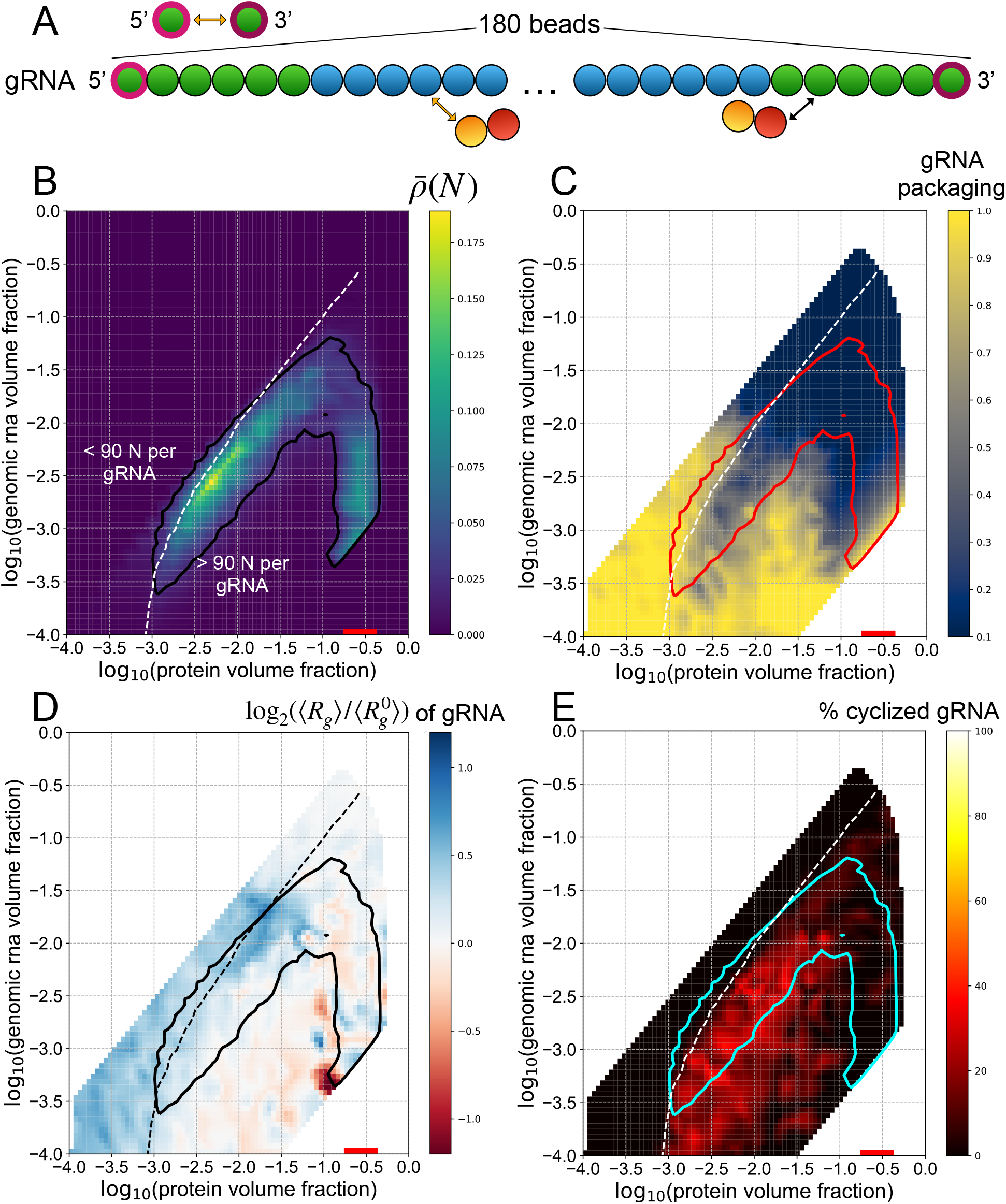
gRNA phase separates with N-protein which limits single genome packaging but promotes genome compaction and cyclization. (A) Genomic RNA (gRNA) is represented as a chain with 180 beads. The terminal 6 beads on the 5ʹ and 3ʹ end are 5ʹ end-like beads, and the rest are FE-like beads. An additional anisotropic interaction among the terminal beads is added to represent known nucleotide complementarity. (B) The phase boundary is drawn along the contour 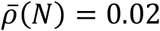 and is shown in black. The heatmap indicates the value of 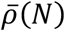 at different volume fractions of N-protein and gRNA. The white dotted line indicates the contour along which an equal volume fraction of N-protein and gRNA is found within a given cluster, i.e. 90 Nproteins for each gRNA. A heatmap indicating the average number of proteins bound to each gRNA is shown in **Fig S10**. This contour is shown in all subsequent panels. (C) The phase boundary is shown in red. The heatmap shows the single gRNA packaging metric, which is the number of clusters containing gRNA divided by the total number of gRNA chains in each simulation. A value of 1 represents perfect single-genome packaging. This metric is shown for an analogous system without attractive interactions in **Fig S9A**. (D) The phase boundary is shown in black. The heatmap shows the fold change of the average radius of gyration of gRNA chains, ⟨*R_g_*⟩, over the average radius of gyration of gRNA in a system without attractive interactions, 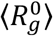. A phase diagram with a heatmap of 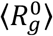 is shown in **Fig S9B**. (E) The phase boundary is shown in cyan. The heatmap shows the percentage of cyclized genomes. Each gRNA chain is categorized as cyclized if its terminal beads are in adjacent lattice positions.

Due to the length of gRNA chains, we were only able to include 10 chains per simulation, with hundreds to thousands of N-proteins. Interestingly, we did not observe dilute phases of gRNA chains under any conditions; there were no well-defined clusters including all gRNA chains, and instead, all gRNA chains were coated with N-protein. N-protein demonstrated a well-defined dilute phase and a dense phase on and around gRNA chains. For these reasons, we determined that phase separation of gRNA in this context was not well-defined, and regarded gRNA instead as a surface upon which N-protein phase separation occurred. To quantify this behavior, we altered the definition of the density inhomogeneity metric, 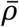, originally defined as a function of the pair distribution functions between all beads in a given simulation as explained in (16). Since we observed a density transition only of N-protein in gRNA-containing systems, we defined a new metric, 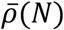 which depends only on the pair distribution function of Nproteins in the system. Thus, the following phase diagrams describing gRNA-containing systems are no longer ternary phase diagrams as in **Figs 1** and **2** with well-defined dilute phases of both components. They instead indicate the density inhomogeneity of N-protein as a function of the volume fraction of N-protein and the surface upon which they condense, gRNA.

For a given N-protein concentration, phase separation of N-protein in the gRNA system occurs over a smaller range of RNA concentrations relative to the 5ʹ end system (**Fig 1C**, **Fig 3B**). This result is consistent with experiments in which 5ʹ end and FE RNAs were combined *in trans* which led to more limited phase separation compared to 5ʹ end RNA alone (11). However, phase separation of N-protein can occur at lower volume fractions of N-protein and gRNA due to the length of the gRNA chains, which is also consistent with experiments that were performed with RNA purified from infected cells that contained gRNA (11). We quantified the average amount of protein bound to single genomes and found that the contour delineating an equal volume fraction of protein and gRNA per cluster aligns well with the high-RNA concentration edge of the phase envelope (white dotted line, **Fig 3B**). The area to the right of this contour indicates bulk concentrations of N-protein and gRNA that lead to a majority of volume fraction per cluster occupied by N-protein. This region includes almost all of the phase separating regime and likely captures the most relevant stoichiometries of gRNA and N-protein during infection in host cells and virion assembly (21).

During virion assembly, single genomes must be packaged within a capsid built of structural proteins and N-protein (22), so we also quantified how many gRNA chains were in each phase separated cluster. We defined a simple metric to quantify single gRNA packaging; the number of gRNA-containing clusters in the system is divided by the total number of gRNA chains. Therefore, the metric is 1 when single-genome packaging is perfect, and approaches 0 as multiple genomes are clustered together. A zeroth-order effect due to excluded volume in the absence of other interactions is shown in **Fig S9A** and demonstrates that single packaging becomes impossible with prohibitively high volume fractions of N-protein or gRNA. In the presence of interactions, our analysis indicates that, for the most part, N-protein phase separation hinders packaging of single gRNAs in clusters (**Fig 3C**). However, at higher N-protein volume fractions in the concave region below the phase envelope, single packaging is robust. It appears that coexisting phases of N-protein condensed on gRNA and a dilute phase in solution promotes clustering of gRNA, but that a more uniform, high density of N-protein throughout the system effectively keeps gRNA chains separated.

If arranged linearly, the ∼30kb genome has an end-to-end length of roughly 10000nm. However, during virion assembly, this genome must be packaged into a viral particle with a diameter of about 100nm, representing an immense compaction challenge. We reasoned that N-protein binding could provide a simple mechanism for gRNA compaction. To quantify compaction, we measured the average radius of gyration of gRNA in each simulation (23),

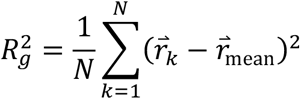

where *N* = 180 is the number of monomers in a chain, 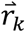 is the position of monomer *k*, and 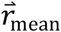 is the average position of monomers in the chain. *R_g_* is computed for each gRNA in a simulation, and the average across gRNAs, ⟨*R_g_*⟩, is reported for each simulation. Again, there is an effect due solely to excluded-volume interactions that leads to compaction of gRNA chains under more crowded conditions (**Fig. S9**). Therefore, we evaluated the effects of N-protein binding interactions using the fold change in the radius of gyration in simulations with interactions relative to those without interactions, 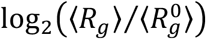. We found that N-protein binding indeed leads to more compact genomes to the right of the equal protein-gRNA volume fraction contour (dotted white line), and more extended genomes to the left of it (**Fig 3D**). As seen with the single genome packaging metric, the most robust compaction occurs in the concave region of the phase diagram with a high N-protein density (**Fig 3D**).

Genome cyclization is important for replication of many RNA viruses (24), and there is recent *in vivo* evidence of cyclization of the SARS-CoV-2 genome (20). We thus sought to characterize the potential role of phase behavior in genome cyclization in our model. We hypothesized that the similar RNA features at the 5ʹ and 3ʹ ends of the genome would promote cyclization when facilitating phase separation of N-protein. We defined a cyclized genome as one whose terminal beads occupy adjacent lattice sites. Averaging over the final half of each simulation, we quantified the percentage of genomes that met this criterion. Strikingly, we see that up to 40-50% of genomes are cyclized for volume fractions to the right of the equal N-protein gRNA volume fraction contour and within the phase boundary (**Fig 3E**). Thus, it appears that sufficient binding of N-protein is essential for cyclization of gRNA molecules, with phase separation providing additional efficiency.

We also performed parameter sweeps for this gRNA-containing system, focusing on the two most sensitive interactions identified in the N-protein with 5ʹ end and FE RNA parameter sweeps; these are the isotropic interaction between N-protein and 5ʹ end RNA and the isotropic interaction between N-proteins. With an increased binding energy between N-protein and the 5ʹ end RNA beads in the gRNA, we saw a modest extension of the phase boundary (**Fig S7A**). Single genome packaging (**Fig S7B**) and compaction (**Fig S7C**) are relatively unaffected with respect to WT, but surprisingly, cyclization is much less efficient (**Fig S7D**). It appears that while N-protein binding to the ends of gRNA chains can facilitate their co-location, very strong binding may limit the ability of the ends of the gRNA chains to come into contact due to interference from N-protein. For the system with increased isotropic interactions among Nproteins, we observed a widening of the phase boundary corresponding to the broader range of volume fractions over which N-protein alone phase separates under this condition (**Fig S8**). Single genome packaging is unaffected (**Fig S8B**), compaction is minimally enhanced in the concave region below the phase envelope (**Fig S8C**), and genome cyclization is unaffected (**Fig S8D**).

### Maximal 5ʹ end-like RNA content in gRNA chains leads to optimal packaging

Our previous work experimentally examined relatively small regions of the genome with regard to the ability to promote N-protein phase separation. While it was clear that the 5’ and 3’ ends of the genome were both highly structured and this was associated with LLPS-promoting activity, it was less definitive that most of the interior of the genome was most similar to FE. Therefore, we next asked how the total LLPS-promoting content vs. the solubilizing RNA content altered N-protein phase separation and genome packaging. We created gRNA chains composed entirely of either 5ʹ end-like beads or FE-like beads (**Fig 4A**). We found that a genome of purely 5ʹ end-like beads led to greatly enhanced phase separation of N-protein (**Fig 4B**), robust gRNA compaction below the equal volume fraction contour (**Fig 4C**), and nearly perfect genome cyclization below the equal volume fraction contour and within the phase envelope (**Fig 4D**). In contrast, gRNA composed entirely of FE beads limited single packaging at high volume fractions of N-protein and gRNA (**Fig 4B**), led to gRNA expansion for most conditions (**Fig 4C**), and did not promote genome cyclization (**Fig 4D**). It appears that maximal content of LLPS-promoting sequence is ideal for packaging gRNA by N-protein alone. However, the genome must encode many features other than optimal binding with N-protein and is thus constrained to have a limited amount of LLPS-promoting sequence. Given a limited supply of LLPS-promoting sequence, we next asked what the optimal spatial patterning would be to promote packaging.

**Figure 4:**
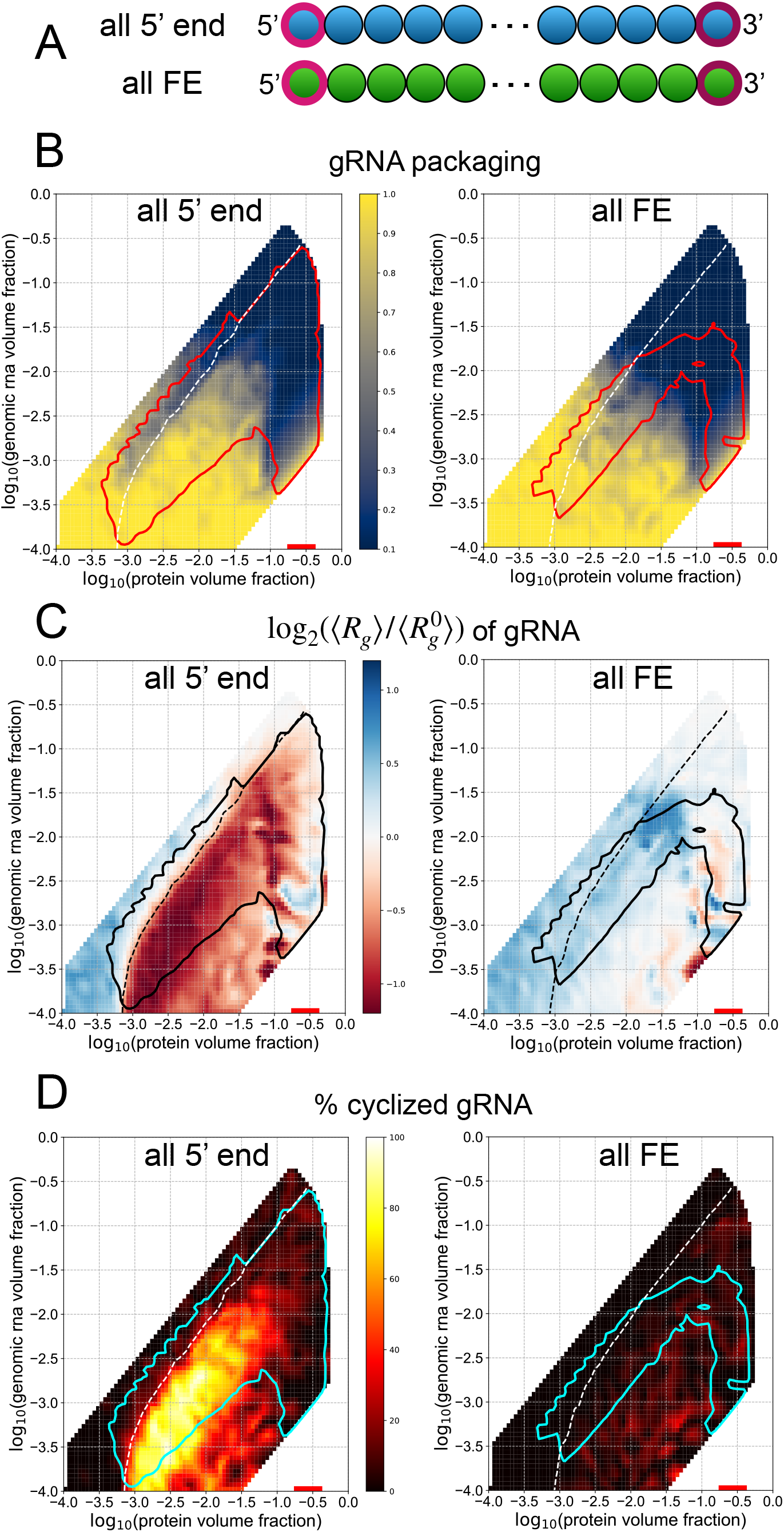
LLPS-promoting sites enhance genome packaging. (A) Schematic showing gRNA chains with only 5ʹ end-like beads and only FE-like beads. (B) The phase boundaries are shown in red. Heatmaps show the single genome packaging metric for each gRNA mutant. (C) The phase boundaries are shown in black. Heatmaps show the fold-change in radius of gyration for each gRNA mutant. (D) Phase boundaries are shown in cyan. Heatmaps show the genome cyclization metric for each gRNA mutant.

### Spatial patterning gRNA mutants can enhance single genome packaging but limit compaction and prevent genome cyclization

Since the 5ʹ and 3ʹ ends of the gRNA were both found to promote phase separation with Nprotein, we hypothesized that this spatial arrangement of phase separation-promoting elements at the ends of the genome may be relevant to packaging. We investigated the importance of the arrangement of phase separation-promoting sequences on the ends of the gRNA by designing mutants where these regions are rearranged. We created three mutants, all of which retain 12 5ʹ end-like beads and the anisotropic interaction among their terminal beads. The 5ʹ end-like beads are repositioned either in the middle of the genome (middle), uniformly throughout (uniform), or on one end (end) (**Fig 5A**). The phase boundaries of each of the mutants remain relatively unchanged with respect to the WT system, suggesting that the total amount and not the spatial patterning of 5ʹ end and FE RNA beads determines the bulk concentrations at which N-protein phase separation occurs (**Fig 5**). However, we observed differences in the genome packaging metrics relative to WT and among the mutants.

The uniform and middle gRNA systems are more efficient than WT at packaging single genomes into clusters, and end gRNA behaves similarly to WT (**Fig 5B**). Thus, it appears that dispersed or centrally located phase separating elements within the gRNA are preferred for single genome packaging by N-protein alone. However, uniform gRNA does not significantly compact upon phase separation, while middle and end gRNA compact similarly to WT, suggesting that sufficiently clustered LLPS-promoting sequences are important for compaction (**Fig 5C**).

**Figure 5:**
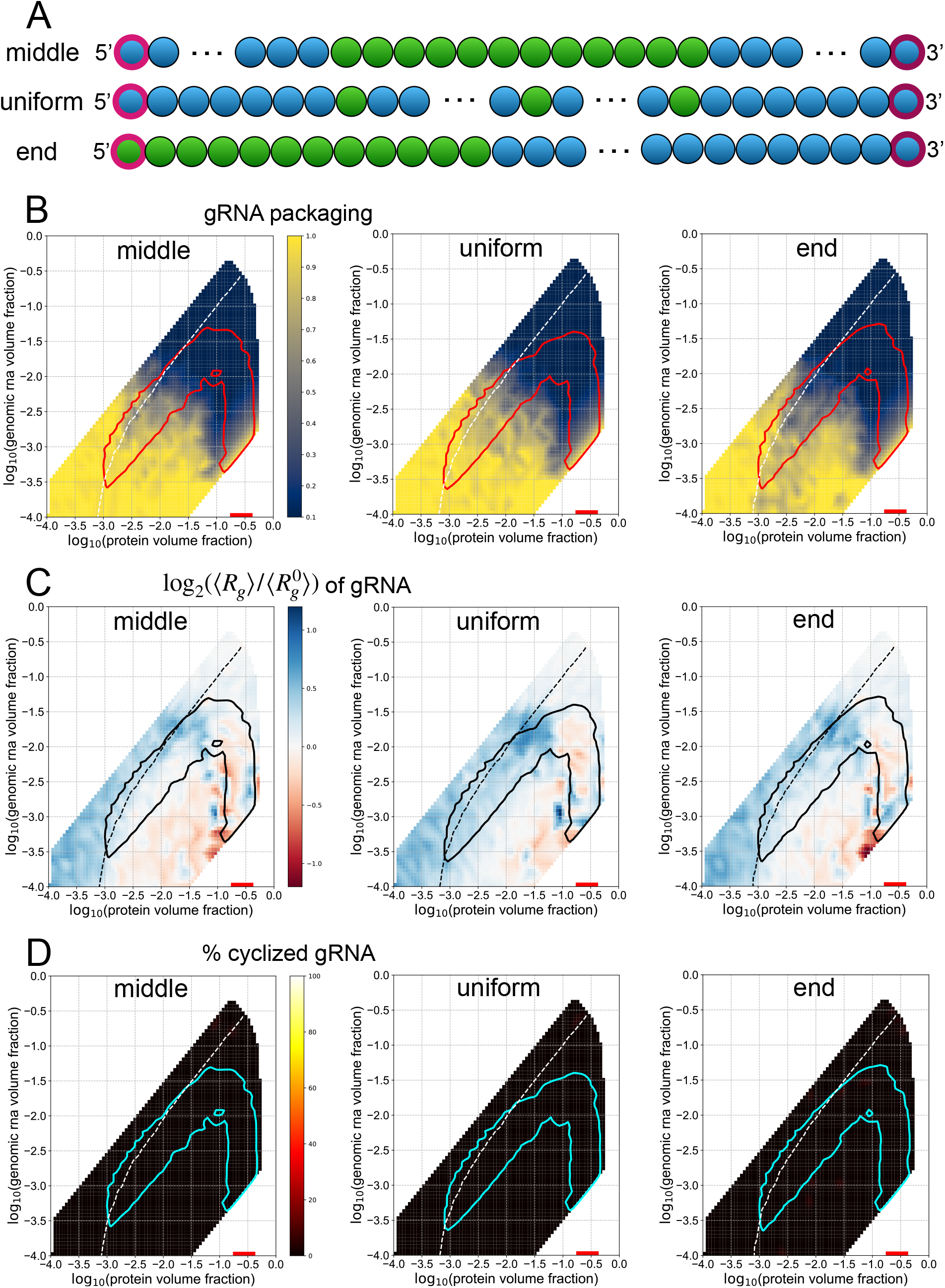
Spatial patterning gRNA mutants show altered genome packaging metrics. (A) Three spatial patterning genome mutants were constructed. Each mutant retains the same number of 5ʹ end-like and FE-like beads as in WT. The terminal beads maintain their anisotropic interaction, regardless of their identity as 5ʹ end-like or FE-like. (B) Phase boundaries are shown in red for each spatial patterning mutant system. The white dotted line indicates the contour along which an equal volume fraction of N-protein and gRNA is found within a given cluster and is included in all subsequent panels. The heatmaps show the single gRNA packaging metric. (C) Phase boundaries are shown in black. The heatmaps show the fold change in the average radius of gyration of gRNA chains. (D) Phase boundaries are shown in cyan. The heatmaps show the percentage of cyclized genomes.

Given the positioning of the wildtype LLPS-promoting sequences at the ends of the genome, we postulated that rearrangement of the location of these sequences would have the strongest impact on genome cyclization. To this end, we also quantified genome cyclization for these mutants, and found that none of them were able to cyclize genomes (**Fig 5D**). Importantly, each of these systems maintains an intrinsic bonding capability between the terminal beads of its gRNA chains. However, since the chains are so large, they cannot efficiently locate each other during the course of the simulation. Thus, the localization of phase separation to the 5ʹ and 3ʹ ends with N-protein in the WT system is necessary for the positioning of the genome ends for binding and cyclization (**Fig 3E**, **5D**).

### Optimal gRNA design with limited LLPS-promoting sequence

We know from the studies above that increased 5ʹ end-like RNA content enhances all packaging metrics. With a limited supply, it appears that positioning of these beads at the ends of genome is essential for cyclization, clustered beads are important for compaction, and uniformly spaced or centrally located beads can promote single packaging. With this understanding, we hypothesized that a genome could evolve that would optimally function according to all of these metrics, given a limited supply of LLPS-promoting beads. We designed a gRNA which has the WT arrangement at the ends, with 12 additional groups of 3 5ʹ end-like beads uniformly spaced throughout its length (Fig 6A). We found that N-protein phase separation occurs over a broader range of concentrations, and that all studied genome packaging metrics are enhanced, relative to WT. Single genome packaging is more preferable within the concave region below the phase envelope (Fig 6B). Genome compaction is greatly enhanced below the equal Nprotein gRNA volume fraction contour (Fig 6C). Genome cyclization is also enhanced (Fig 6D).

**Figure 6:**
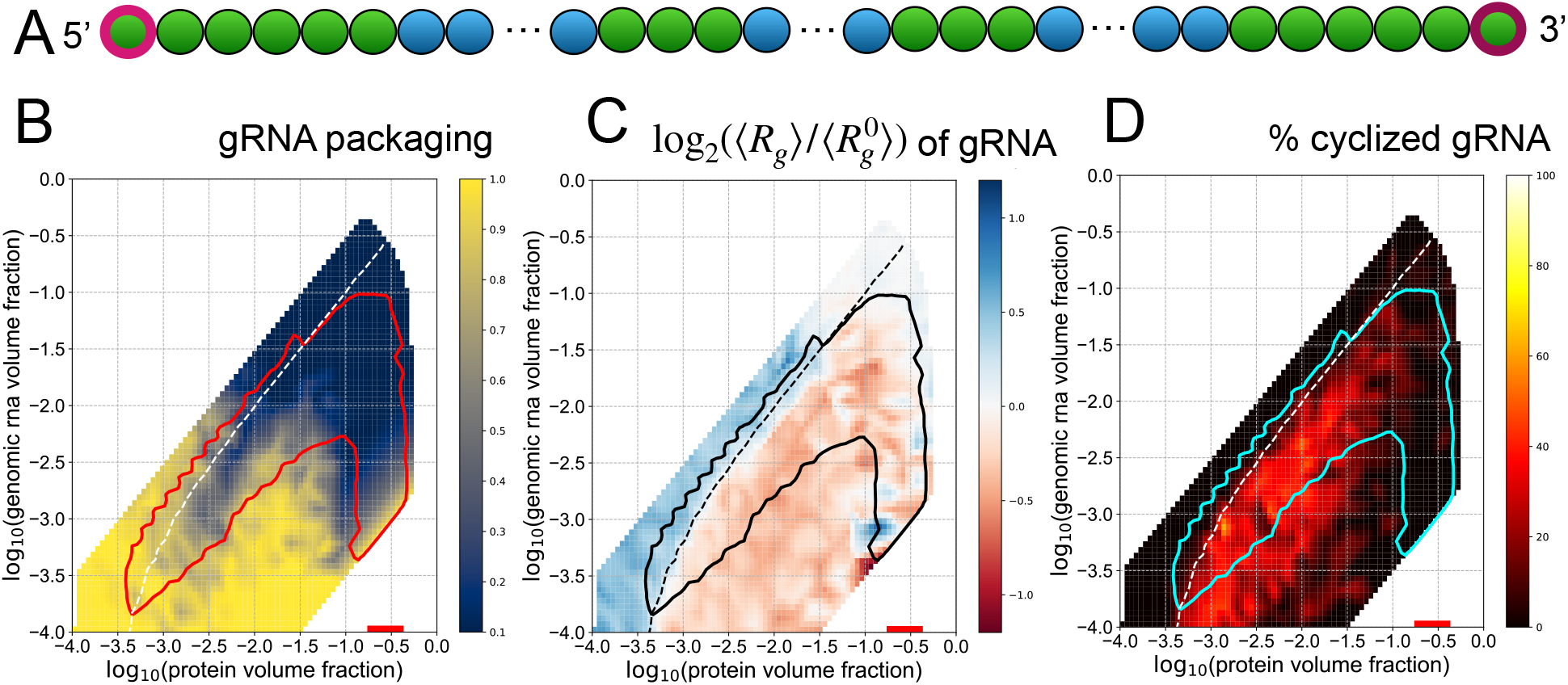
WT and uniform elements optimize genome packaging when combined. (A) The optimal gRNA pattern for singular and compact packaging has 6 5ʹ end-like beads at each end, with 12 additional groups of 3 5ʹ end-like beads distributed uniformly throughout the rest of the chain. (B) The phase boundary is shown in red. The white dotted line indicates the contour along which an equal volume fraction of N-protein and gRNA is found within a given cluster and is included in all subsequent panels. The heatmap shows the single gRNA packaging metric. (C) The phase boundary is shown in black. The heatmap shows the fold change of the average radius of gyration of gRNA chains relative to a system with only excluded volume interactions. (D) The phase boundary is shown in cyan. The heatmap shows the percentage of cyclized genomes.

For this optimized gRNA design, there appears to be a concentration regime corresponding to the concave region below the phase envelope where single genome packaging, compaction, and cyclization can all occur efficiently. These results raise two predictions concerning the virion assembly process. First, there exists an optimal concentration range of gRNA and N-protein that promotes virion assembly. Second, phase-separation promoting gRNA sequences may be located not only at the 5ʹ and 3ʹ ends, but also distributed in clusters throughout the genome to enhance genome packaging.

## Discussion

It is clear that protein and RNA elements of SARS-CoV-2 can engage in phase separation with Nprotein (11–15), but the functional consequences of this physical chemistry capacity for viral replication remain elusive. In this study we sought to explore how the spatial patterning of phase separation-promoting or -inhibiting RNA elements in the genome could facilitate the specificity and singularity of packaging the genome. Using coarse-grained simulations rooted in empirical observations, we find that single genome packaging is most efficient when binding sites are centrally located or distributed throughout the genome. However, the arrangement of phase separation promoting sequences in clusters is critical for genome compaction, and the positioning of these elements at both ends is necessary for cyclization.

### Biophysical interpretation of the different protein binding modes leading to opposing phase behavior

An essential feature of our model is the different binding mode of N-protein with 5ʹ end and FE RNA. We found that weak, isotropic interactions with 5ʹ end RNA promote phase separation, while strong, anisotropic interactions with FE RNA lead to N-protein solubilization. Intrinsically, the valence of the isotropic interactions is 26 in a cubic lattice, while the valence is 1 for the anisotropic interactions. Additionally, since the anisotropic interactions have a higher binding energy than the isotropic ones, they last for longer, which effectively compounds the difference in valence between the 5ʹ end and FE beads. For the FE RNA system, this design allows for more N-protein binding at lower protein concentrations, but a less significant increase in protein binding as the protein concentration is increased. On the other hand, 5ʹ end RNA experiences a cooperative binding effect with N-protein, leading to greatly increased protein binding as protein concentrations are increased. The valence difference between 5ʹ end and FE RNA also includes competition between N-protein-FE RNA binding and N-protein dimerization, since a single bead in the model is responsible for both interactions. Therefore, while N-protein is bound to FE RNA, it can no longer dimerize, but while it is bound to 5ʹ end RNA, it is free to dimerize. We suspect that this competition also compounds the cooperative effect of the Nprotein binding to each of these different RNAs and contributes to the distinct behaviors of these different polymers. The agreement between experimental results and our model suggests that there may be some underlying differences in the N-protein-RNA interactions in the 5ʹ endlike and FE-like regions in the gRNA that lead to distinct protein binding behavior. We hypothesize that FE-like regions have different N-protein binding kinetics than 5ʹ end-like regions, which will need to be explored experimentally in future work.

### Boomerang shape of the phase boundaries

It has been previously reported that phase boundaries in ternary systems that are purely driven by heterotypic interactions are roughly elliptical in log-log space (16). However, the phase diagrams of the ternary systems studied in this work have distinct shapes. The 5ʹ end RNA with N-protein phase diagram resembles an elliptical shape combined with a high N-protein concentration arm that corresponds well to the shape of the FE RNA with N-protein phase diagram. Each of these high N-protein concentration arms correspond to the N-protein concentration range where phase separation occurs for N-protein alone at temperature 1 a.u., the temperature at which all ternary systems were studied (**Fig 1B-D**). The same superimposed shape is even more apparent in the gRNA with N-protein phase diagrams, with N-protein phase boundaries that resemble boomerangs (**Fig 3B-E**). We hypothesize that these phase diagrams arise from the union of two regimes of phase behavior that are driven by distinct forces. The elliptical portion that extends to low N-protein and RNA concentrations demonstrates phase separation that is driven by heterotypic interactions between RNA and N-protein, which aligns with reports in (16). On the other hand, the high concentration N-protein portion of the phase diagrams indicates phase separation that is driven by N-protein homotypic interactions. Here, RNA can partition into the dense phase, but it is neither necessary for phase separation, nor is it the driver. Therefore, the FE RNA with N-protein phase diagram consists only of this high Nprotein concentration regime, which is eventually capped at high enough RNA concentrations (**Fig 1D**). As more FE RNA is present to sequester N-protein out of solution, the concentration of the available pool of N-protein is effectively decreased and phase separation can no longer occur. For systems with 5ʹ end RNA present, the phase diagrams have a complex reentrant character, passing in and out of the phase separating regime for certain fixed RNA concentrations as N-protein concentration is changed (**Fig 1C**, **3B-E**). We speculate that this reentrant behavior is due to the interplay of phase separation driven by heterotypic or homotypic interactions, with intermediate regimes where neither are strong enough to drive phase separation. It is interesting to speculate whether such rich phase behavior may exist in other multicomponent systems, specifically those that involve long nucleotide polymers and RNA- or DNA-binding proteins.

### Compatibility with the single packaging signal model

A recent model of single genome packaging has emerged as part of a study of the SARS-CoV-2 N-protein (13). Results from Cubuk et al. suggests a single packaging signal is much more efficiently packaged and multiple viruses such as HIV employ this strategy. Our results confirm this finding in that restriction of LLPS promoting elements to the ends of the genome (Fig 5) is also not as efficient as single central packaging signal and multiple peppered LLPS promoting elements in the center (**Fig 6**). Cubuk et al showed that for a 2-bead representation of N-protein that experienced isotropic attraction to itself and a long, 61-bead RNA molecule, large phase-separated clusters would form. However, if a much stronger binding site was added to the center of the RNA chain, mimicking a hypothetical packaging signal, N-protein and RNA chains would instead form kinetically trapped clusters that only very slowly coalesced into a single phase-separated droplet. This model is simpler than the one presented here, but it is reminiscent of the distinct effects seen here between 5ʹ end RNA and FE RNA. In our model, the most important effective difference between the 5ʹ end and FE RNA beads is their valence, with 5ʹ end RNA having a much higher valence than FE RNA. The model in (13) includes only a difference in binding energy, since all beads interact isotropically. However, the higher binding energy beads have a lower effective valence than the low energy beads at a given timescale, since they participate in bonds with fewer partners. In line with our results, the presence of lower-valence binding sites sequesters N-protein into clusters, opposing large-scale phase separation. Indeed, other groups have shown that a high enough valence is required for phase separation to occur *in silico* (25) and *in vitro* (26). In our model, however, most of the binding sites on the gRNA are of a low-valence character, which is distinct from the single, low-valence, packaging signal site in (13). Despite this difference, both models provide evidence that such low-valence sites are essential for packaging tasks required during virion assembly, and that runaway phase separation must be tempered via alternative self-assembly pathways.

Why would SARS-CoV-2 and other betacoronaviruses use a relatively inefficient packaging methodology with LLPS? One possible reason could be that packaging and condensation may be acting in direct competition with other viral processes such as translation, and N-protein LLPS may block or slow ribosomal read-through. In line with this hypothesis, Fmr1 LLPS has previously been demonstrated to repress translation of co-condensing RNA (27). By restricting LLPS promoting elements to the 5’ and 3’ ends of the genome, SARS-CoV-2 could allow for efficient packaging while ensuring viral protein production can proceed unencumbered.

### The phase behavior of systems with very short and very long chains

For our systems with N-protein and gRNA, we never observed a dilute phase of gRNA and instead saw that the dense phase of N-protein was condensed on gRNA chains. We hypothesize that this effect is due to the large size of gRNA and their low abundance (10 chains in simulations). The Flory-Huggins free energy (28) predicts that as polymer length increases, the dilute phase volume fraction of that polymer decreases. As chain length increases in a fixed volume, the expected dilute phase volume fraction will eventually pass below the volume fraction occupied by a single chain, leading to disappearance of the dilute phase. When a small number of such large chains are combined with many thousands of shorter chains, an interesting blend of properties can arise. The shorter chain may still exhibit thermodynamically well-defined phase separation, although the dense phase now occurs on the long chains which effectively become a surface upon which phase separation is favored. This situation may be a relatively common one in biology. For example, within a host cell, the number of SARS-CoV-2 gRNA chains in a volume relevant for N-protein phase separation may not be much more than the 10 studied here (9). The difference in scale between thousands of proteins and dozens of very large nucleic acid polymers presents challenges to existing physical frameworks for ternary phase separation and will require theoretical innovations to rigorously understand.

### Generalization to other viruses and systems with long RNAs or DNA

Our models developed here are sufficiently coarse-grained to speculate that they may be applicable to other viruses and systems that involve long nucleotide chains and proteins. Components from several viruses have been shown to undergo phase separation, raising the possibility that spatial patterning of specific LLPS-promoting RNA or DNA sequences may have evolved to promote optimal genome packaging in other viruses in addition to SARS-CoV-2.

Many cellular phase separated bodies involve long RNAs or DNA and proteins that bind them. A particularly relevant example for our modeling is paraspeckles. Paraspeckles are highly ordered, condensed nuclear bodies that require the presence of the long noncoding RNA, NEAT1. NEAT1 was recently shown to contain distinct functional domains, one of which is repetitive in its sequence and is necessary for paraspeckle formation (29). Paraspeckles also require several proteins, most of which contain RNA-binding domains and disordered regions (30). Recent work has shown that the central region of NEAT1 is necessary and sufficient for paraspeckle formation, and that it initiates assembly by binding several proteins (29). Specifically, the proteins NONO and SPFQ must first bind NEAT1, dimerize, and promote further polymerization via coiled-coil domains with other proteins for paraspeckle assembly to continue (30). There are thus many parallels with the SARS-CoV-2 system studied here; specific spatial patterning of protein binding elements along the RNAs is essential, and the protein partners must be able to dimerize/oligomerize for further assembly.

## Conclusion

Identification of specific RNA sequences that promote ordered phase separated bodies via protein binding will not only advance our understanding of viruses, but also the many diverse cellular bodies and regions that contain long RNAs or DNA.

## Author Contributions

S., A. S. G., and C. A. R. designed the conceptual models; I. S. implemented and performed simulations, data analysis, and created figures; I. S., A. S. G., and C. A. R. wrote the manuscript.

## Acknowledgements

We thank Mikayla Feldbauer for helpful discussions and code drafting and the Pappu lab for initial introductions to working with LASSI, and Furqan Dar for technical guidance when modifying LASSI. This work used the Extreme Science and Engineering Discovery Environment (XSEDE), which is supported by National Science Foundation grant number ACI-1548562. The Comet cluster at the San Diego Supercomputer Center was used through allocation nca106. We thank UNC Chapel Hill campus champion Mark Reed for helping I.S. access XSEDE resources on short notice. This work was supported by a FastGrant to ASG and C.A.R. is supported by NIH T32 CA 9156-43, F32GM136164, and L'OREAL USA for Women in Science Fellowship, and I. S. and A. S. G. were supported by NIH R01 GM081506 and the Air Force Office of Scientific Research (grant FA9550-20-1-0241). A. S. G. serves on the scientific advisory board to Dewpoint Therapeutics. The authors declare no conflicts of interest.

