## Supplemental Information for "Role of spatial patterning of N-protein interactions in SARS-CoV-2 genome packaging"

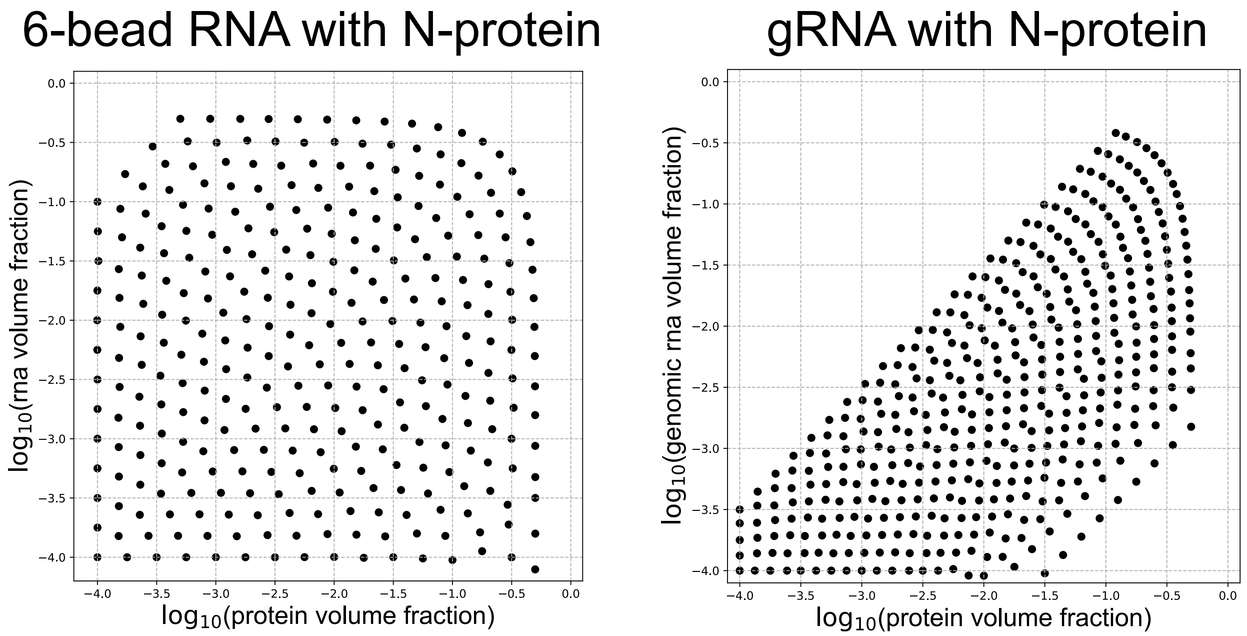


**Figure S1: (protein, gRNA) volume fractions used in ternary simulations.** 4 simulations were run at each point for respective systems as described in methods. Specifically, a given point represents a certain volume fraction of both the protein and RNA components in a system. The total number of chains for each point is set as specified in the methods, and the number of each protein and RNA chain is set to satisfy the corresponding volume fractions at each point.

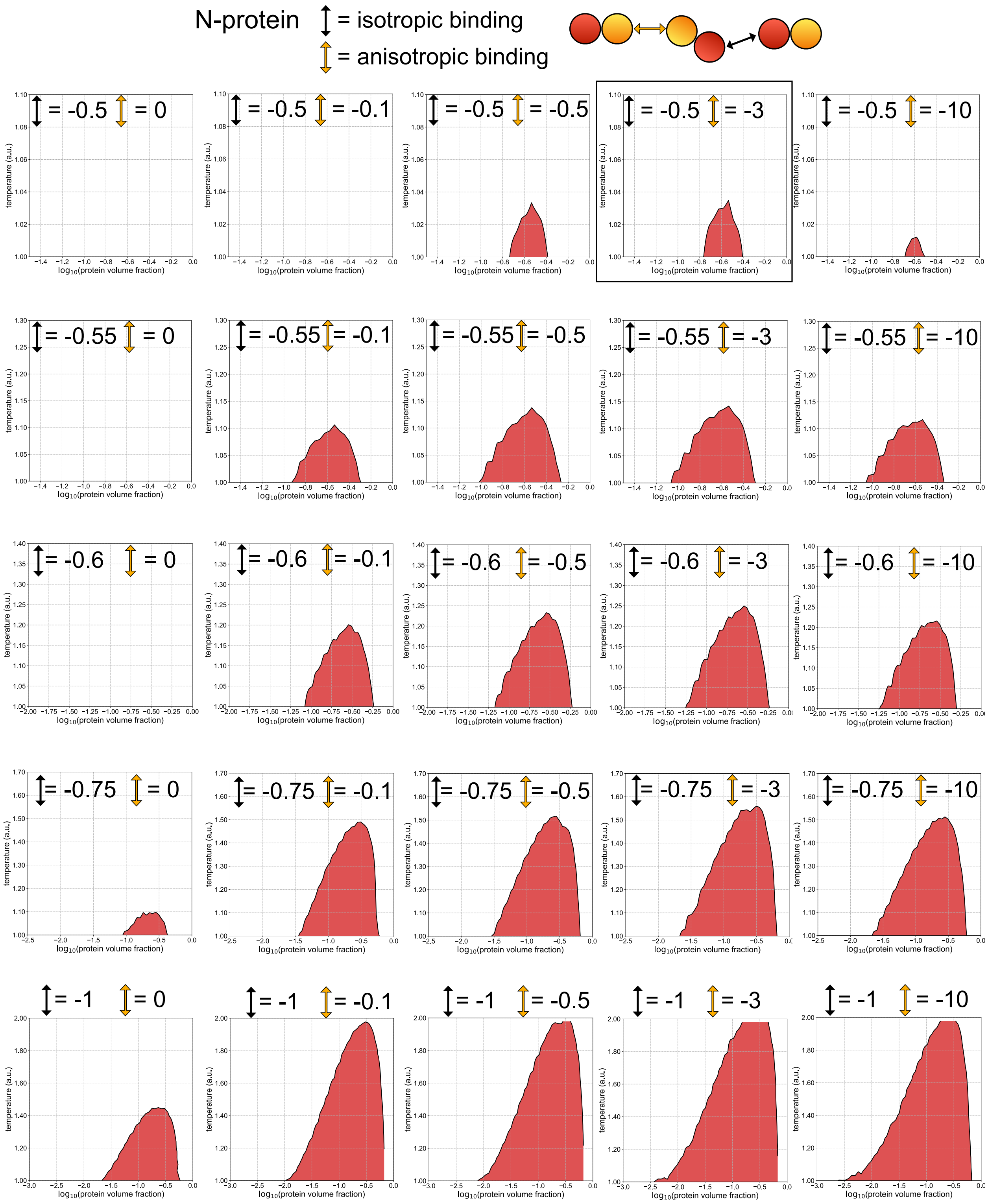


**Figure S2: Sweeps over protein-protein interaction energies for the protein-only system.** An independent set of simulations was performed to generate phase diagrams in each panel shown. Along rows, the isotropic energy is fixed and the anisotropic energy changes, and across columns the anisotropic energy is fixed and the isotropic energy changes. The default N-protein energies are indicated by the black rectangle.

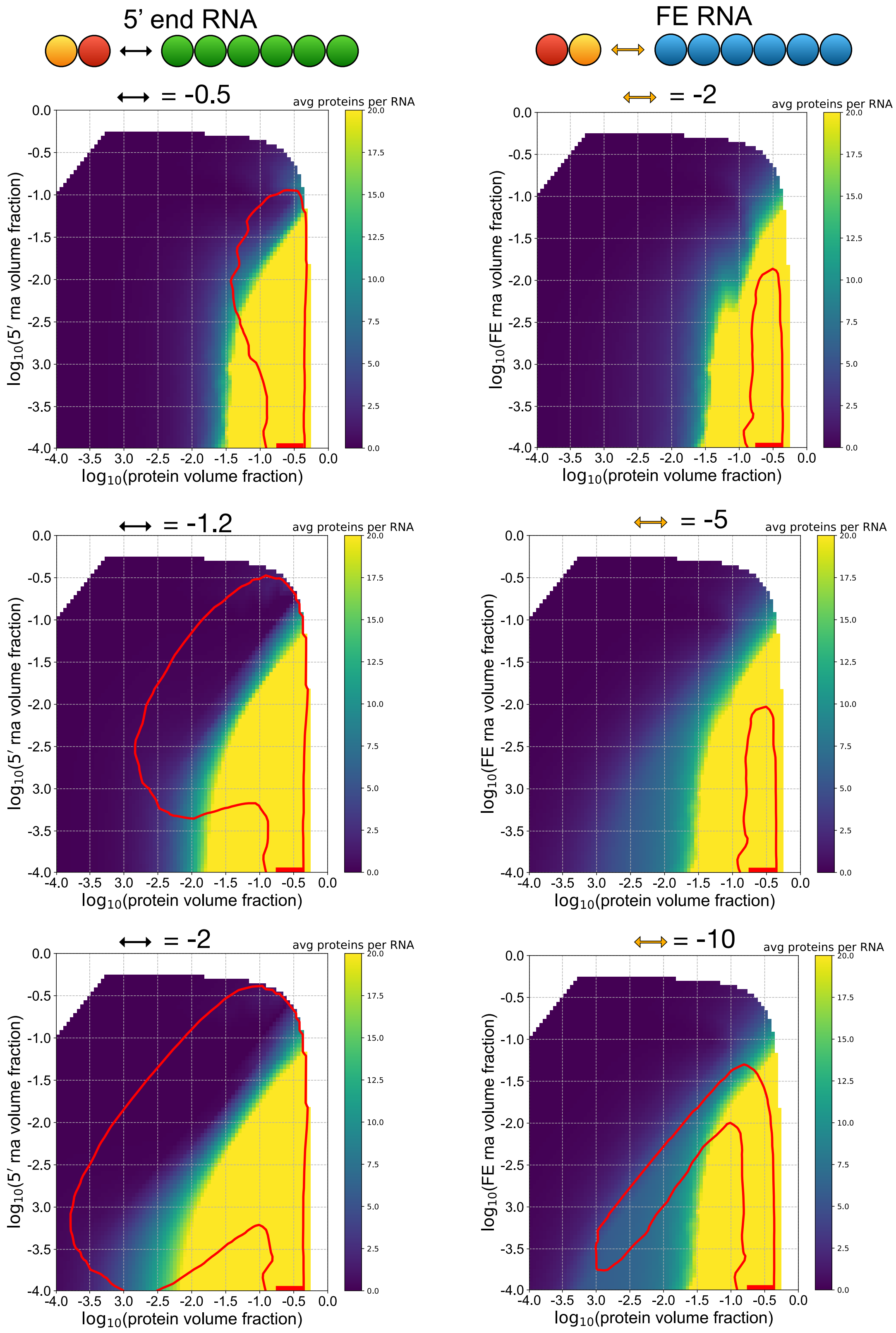


**Figure S3: Sweeps over interaction energies between N-protein and 5’ end RNA and N-protein and FE RNA.** For default N-protein binding energies, the isotropic binding between N-protein and 5’ end RNA is changed along the first column, and the anisotropic binding between N-protein and FE RNA is changed along the second column.

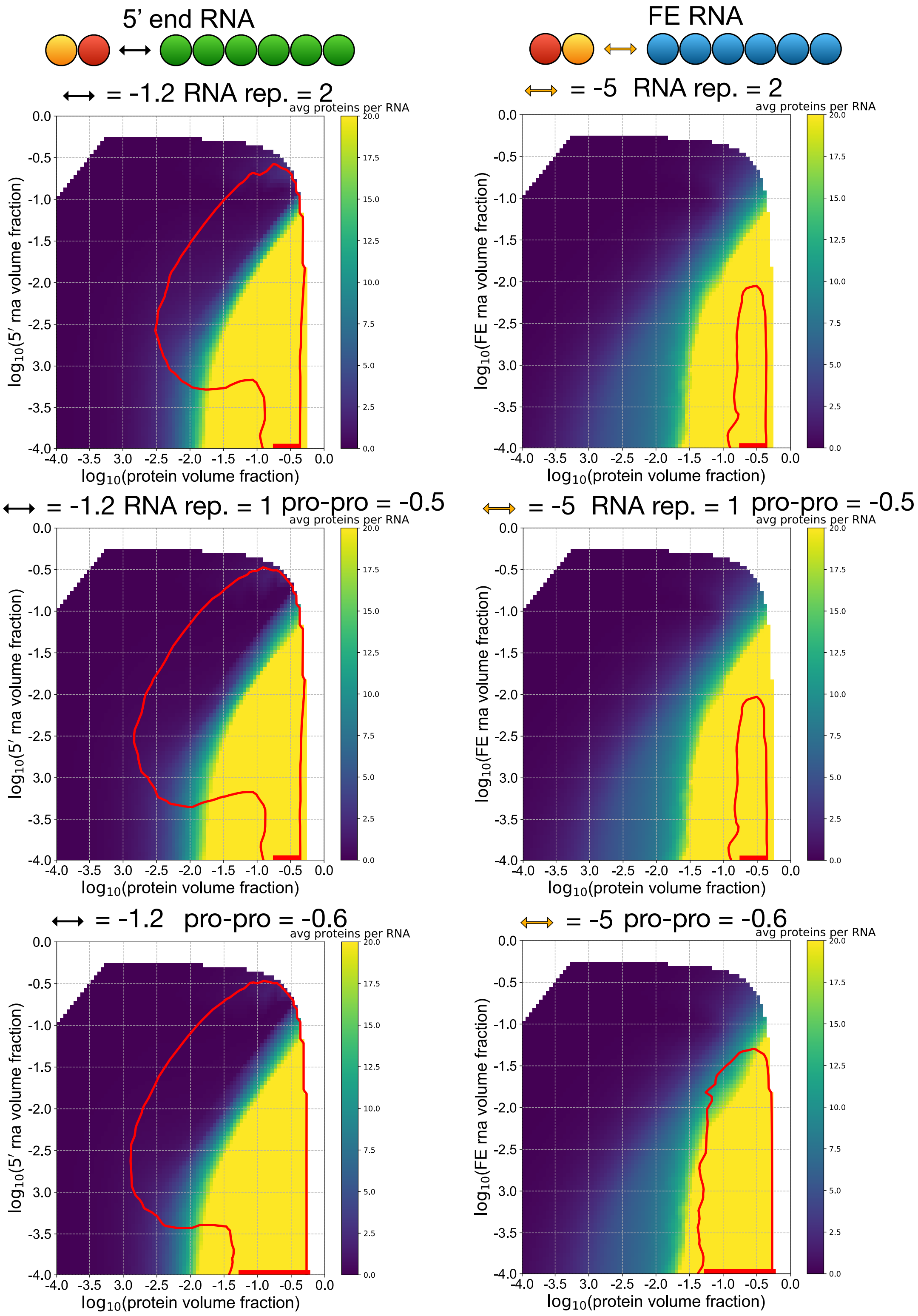


**Figure S4: Results for altered RNA-RNA repulsion energies and protein-protein isotropic interaction energies.** The first row demonstrates the effect of a doubled RNA-RNA isotropic repulsion, with all other parameters set to default values. The second row shows results for default values of all parameters. The third row shows the effect of incorporating N-protein with a higher isotropic self-binding energy.

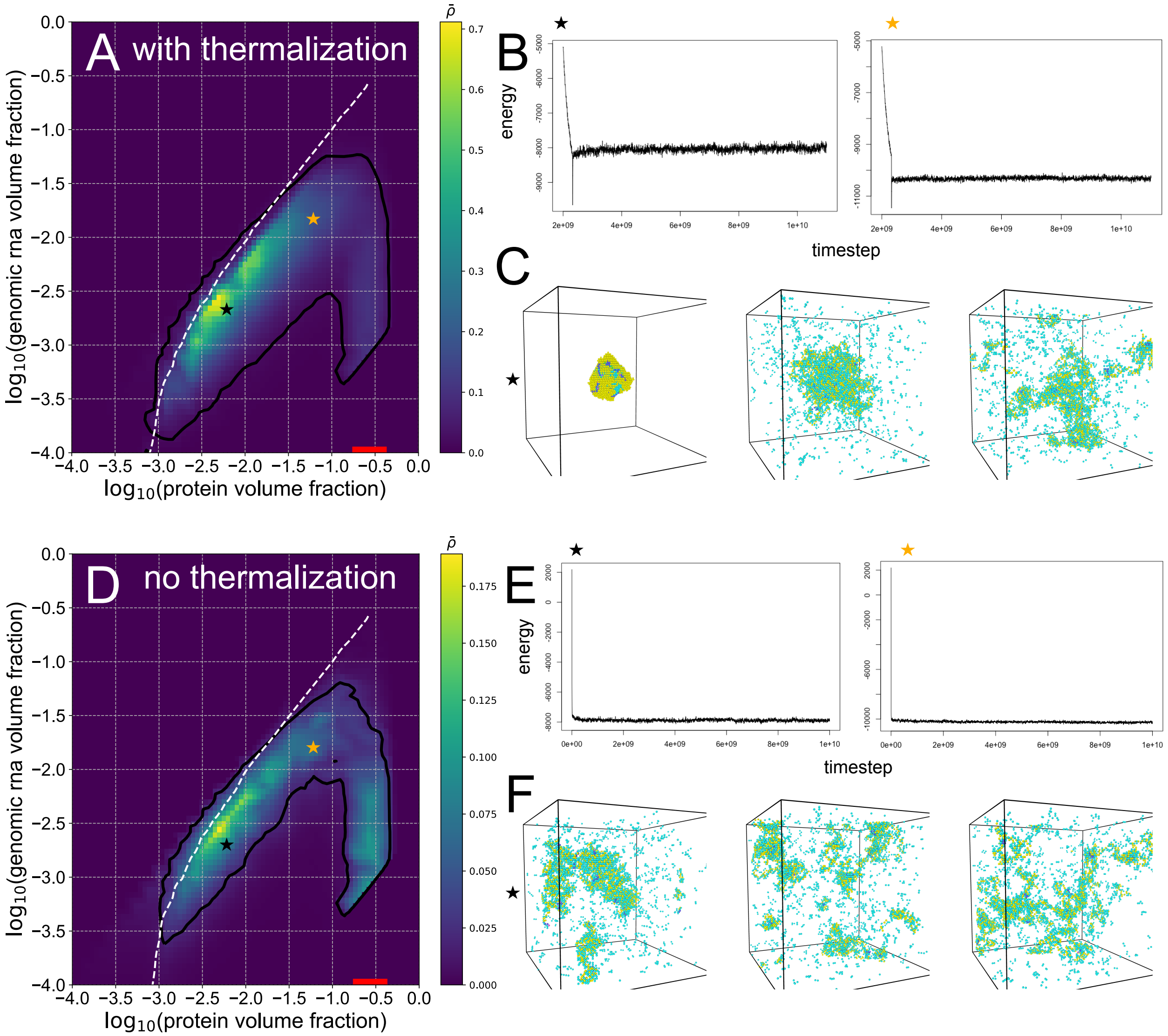


**Figure S5: Comparison of energies and system conformational states with and without thermalization for the default N-protein-gRNA parameters.** (A) Phase diagram for a system run with 1e9 steps of thermalization. (B) Total energy versus timestep plots for two simulations corresponding to black and orange stars in (A). (C) Snapshots of the black star simulation at the beginning during simulation, halfway through, and at the end of the simulation. (D) Phase diagram for the system run with no thermalization. (E) Total energy versus timestep plots for two simulations corresponding to black and orange stars in (D). (F) Snapshots of the black star simulation at the beginning of the simulation, halfway through, and at the end of the simulation.

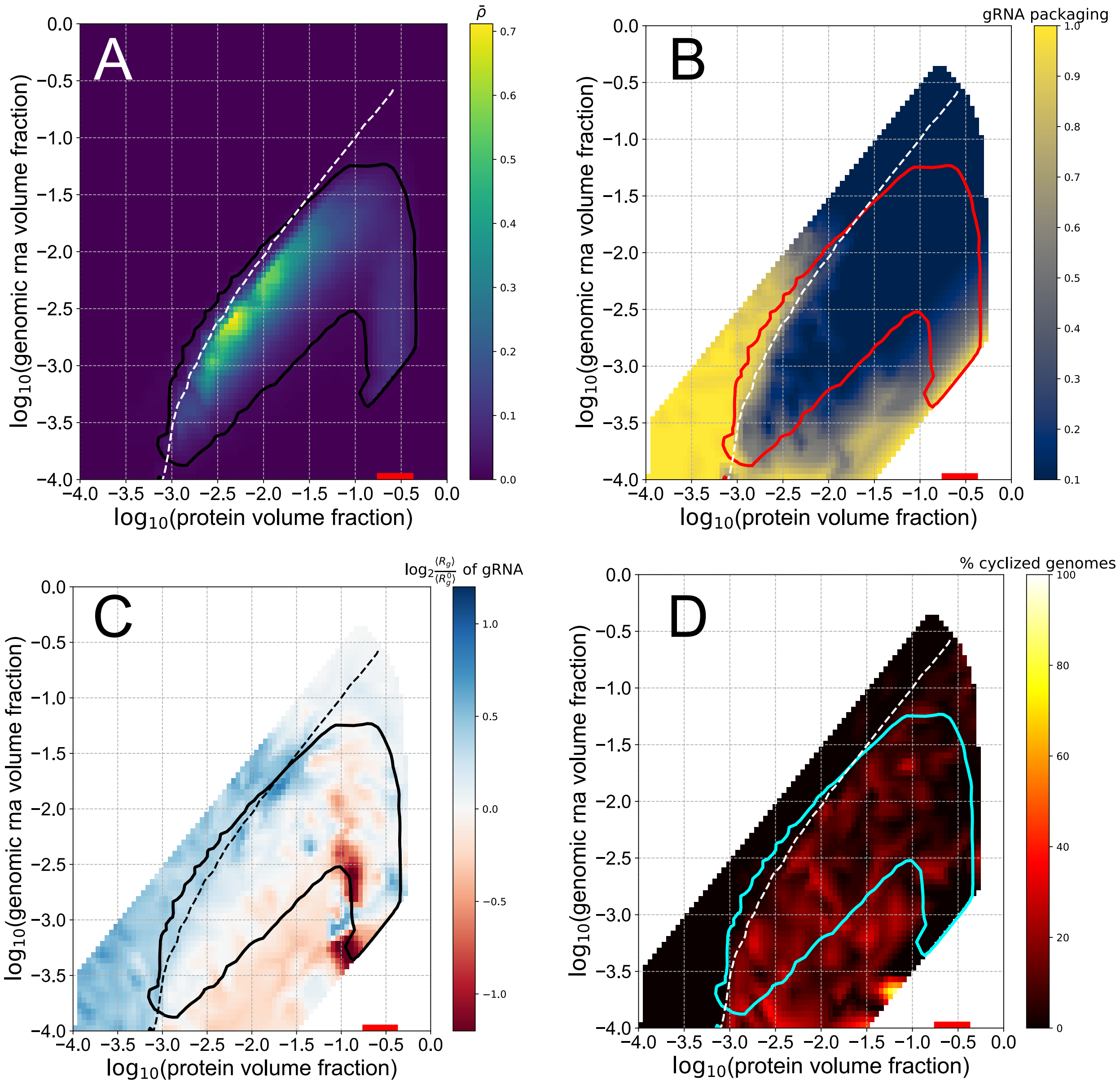


**Figure S6: Phase behavior of the default N-protein with gRNA system with 1e9 steps thermalization at beginning of simulation.** (A) Heatmap shows $\bar{\rho}$ calculated for N-protein – N-protein interactions only, as described in methods. The black line is the contour drawn at $\bar{\rho} = 0.02$, and the white line denotes the bulk volume fractions at which an equal volume fraction of protein is bound to each gRNA molecule, on average. (B) Phase envelope in red overlaid on the single packaging metric. (C) Phase envelope in black overlayed on the fold change in the radius of gyration relative to a system with no interactions. (D) Phase envelope in cyan overlayed on the percentage of cyclized genomes.

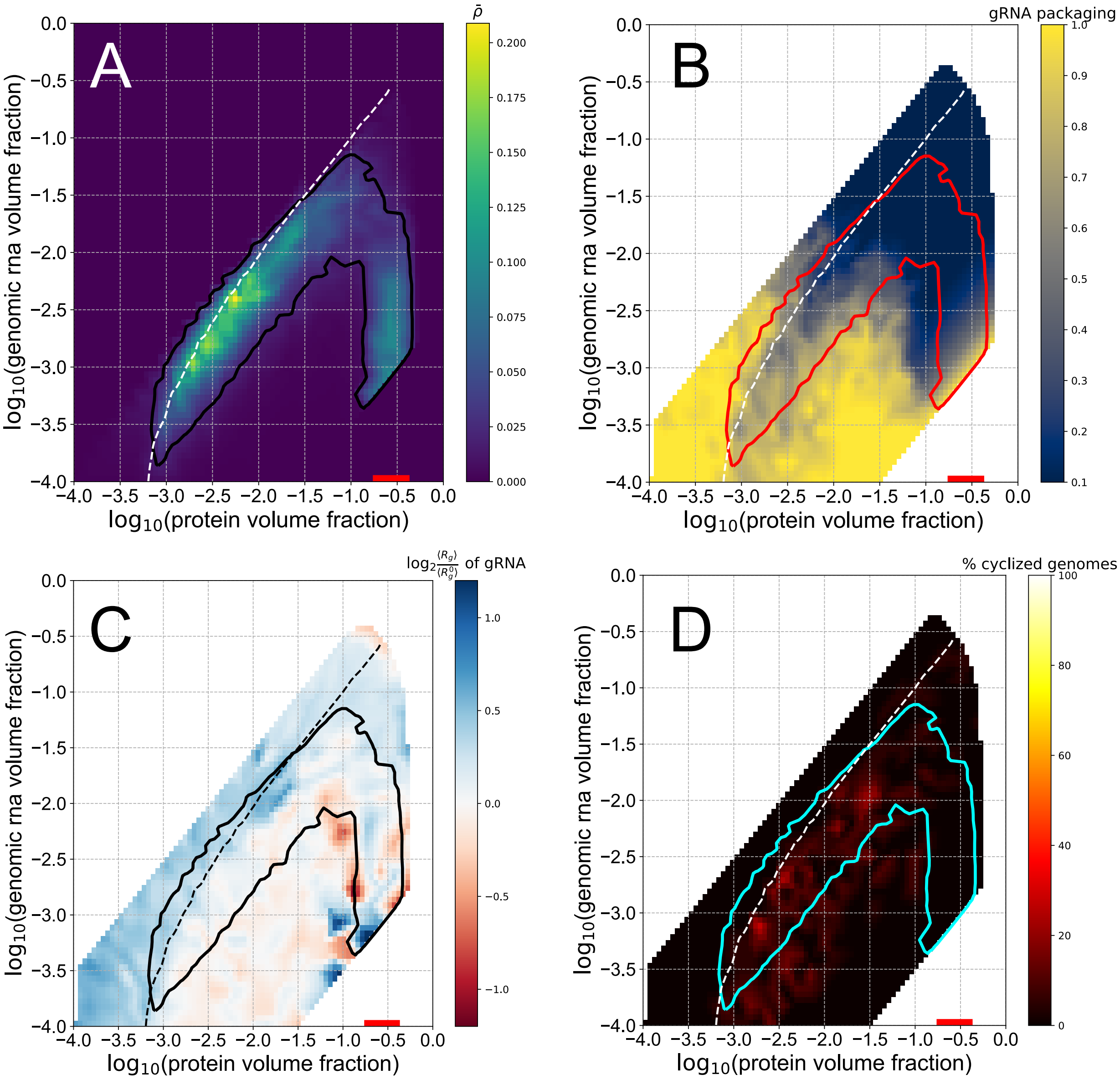


**Figure S7: Phase behavior of N-protein with gRNA with isotropic binding energy between N-protein and 5′ end RNA in the gRNA increased to -2.** (A)-(D) Related to panels in **Fig. S6**.

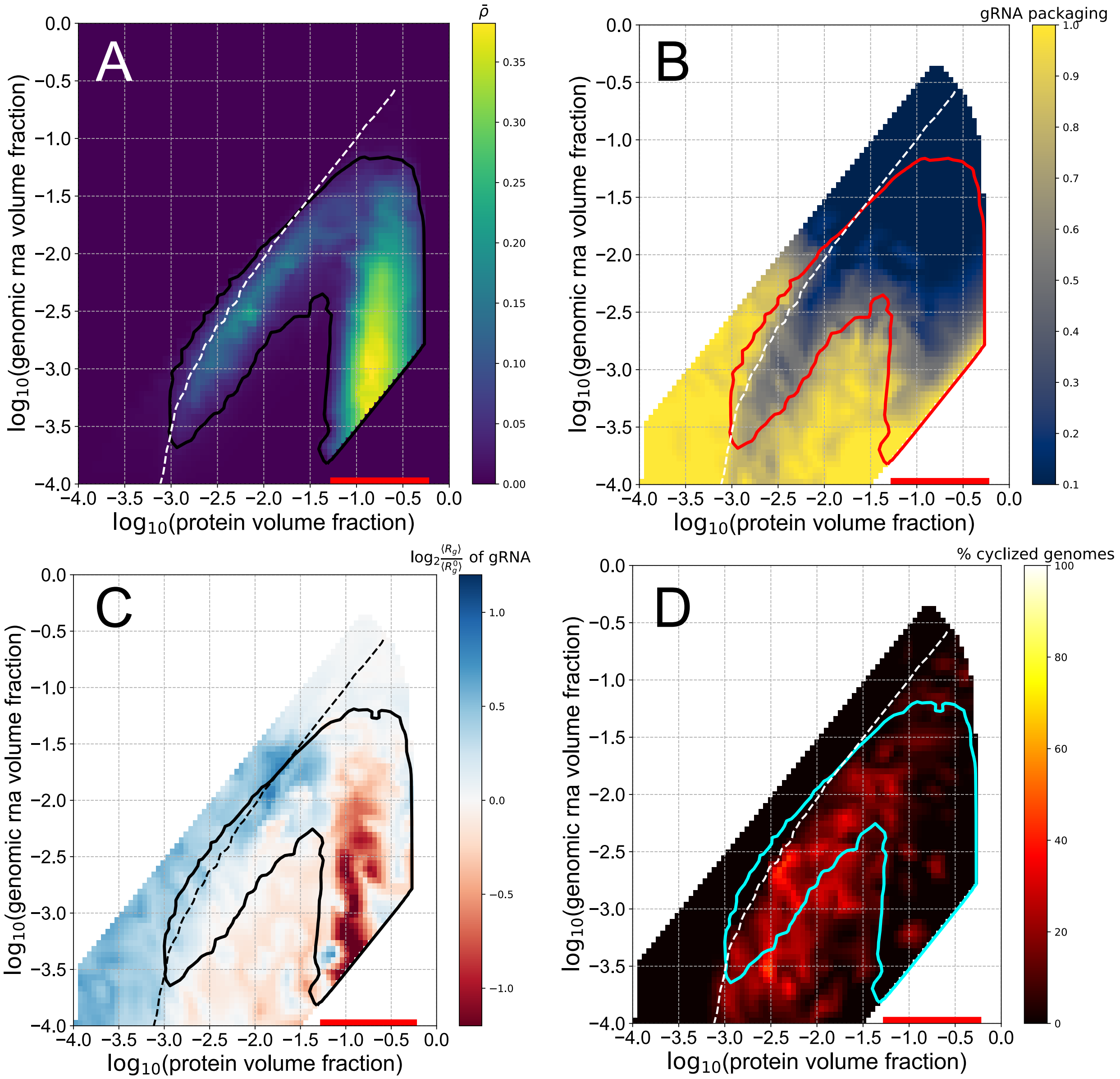


**Figure S8: Phase behavior of the N-protein with gRNA system with N-protein – N-protein isotropic interactions increased to -0.6.** (A)-(D) Related to panels in **Fig. S6**.

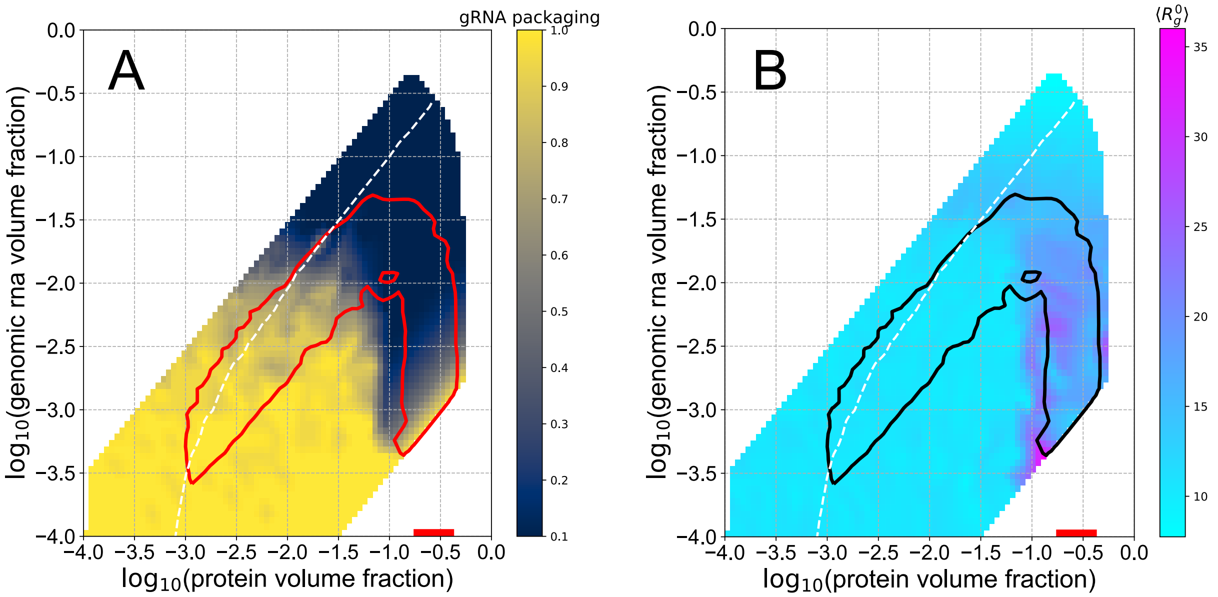


**Figure S9: gRNA packaging and radius of gyration for simulations with no interaction energies.** The only interactions among particles are excluded volume such that only one bead can occupy a lattice site at a given time.

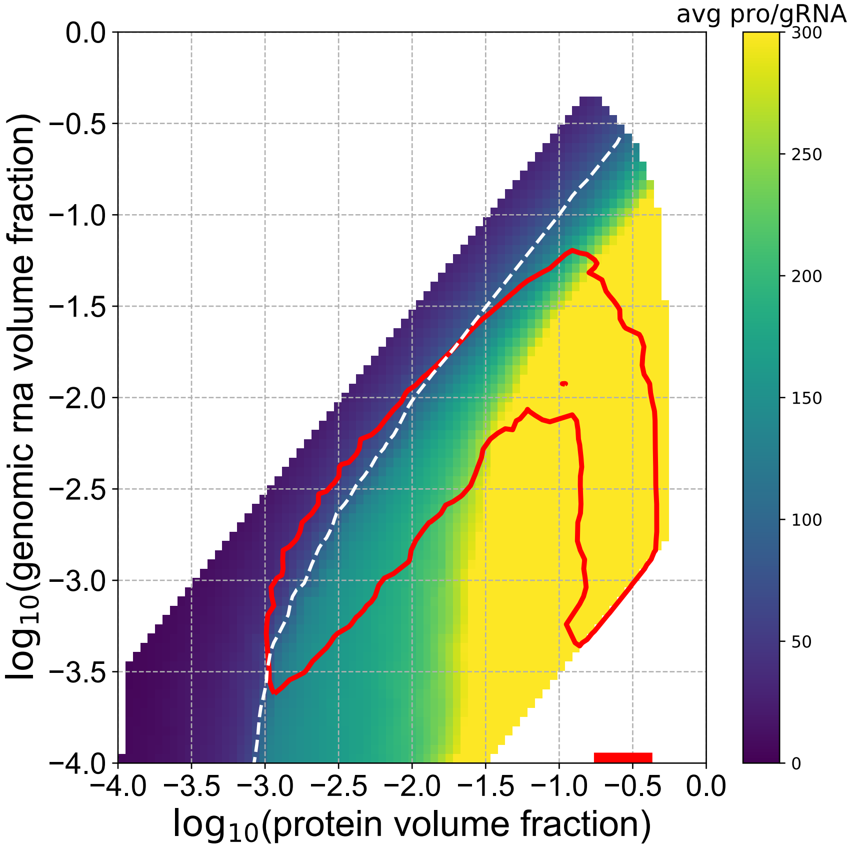


**Figure S10: Average number of N-proteins bound to each gRNA for the WT system.** Phase envelope is drawn in red. The heatmaps indicate the average number of protein chains per gRNA chain for each cluster identified in each simulation. If more than one gRNA chain is in a cluster, the ratio of protein chains to gRNA chains within that cluster is reported.
